# The porcine cGAS-STING pathway exerts an unusual antiviral function independent of IFN and autophagy

**DOI:** 10.1101/2022.07.28.501948

**Authors:** Sen Jiang, Nengwen Xia, Jia Luo, Youwen Zhang, Qi Cao, Jiajia Zhang, Yuening Wang, Wanglong Zheng, Nanhua Chen, François Meurens, Jianzhong Zhu

**Author notes:** Address correspondence and reprint requests to Dr. Jianzhong Zhu, Suite 404, International Cooperation Building, Wenhui Campus, Yangzhou University, 120 University Drive, Yangzhou, 225009, China.

## Abstract

The innate immune DNA sensing cGAS-STING pathway exerts strong antiviral activity through the downstream interferon (IFN) production; however, it has been recently recognized that IFN independent activity of STING also plays an important role in antiviral functions. Nevertheless, the IFN independent antiviral activity of STING is not fully understood. In this study, we showed that porcine STING (pSTING) played a critical role in anti-HSV1 and anti-VSV infections, and IFN defective mutants including pSTING pLxIS sub, S365A and ΔCTT all exhibited similar antiviral functions to wild type (WT) pSTING. Further, all these IFN defective pSTING mutants possessed a comparable autophagy activity relative to WT pSTING as expected. From pSTING WT, S365A and ΔCTT, the residues responsible for autophagy were mutated, which included L333A/R334A, Y167A/L170A and Y245A/L248A, respectively. Surprisingly, all these autophagy defective pSTING mutants still resisted from the two viral infections, demonstrating the pSTING antiviral function independent of IFN as well as autophagy. On the other hand, all the autophagy defective pSTING mutants triggered cell apoptosis, which was associated with the antiviral functions. Additionally, pSTING lost its antiviral activity in TBK1^-/-^ and IRF3^-/-^ porcine macrophages, indicating the involvement of TBK1 and IRF3 in other STING activity such as apoptosis. Collectively, our results revealed that STING exerts both IFN and autophagy independent antiviral activity, and also suggested that STING triggered cell apoptosis might resist from virus infections.

## Introduction

Innate immunity is the first line of defense against pathogenic microorganisms, recognizing the pathogen associated molecular patterns (PAMPs) and damage associated molecular patterns (DAMPs) (1, 2). The innate immune encoded pattern recognition receptors (PRRs) are responsible for recognitions of PAMPs and DAMPs, encompassing membrane bound Toll-like receptors (TLRs) and C-type lectin-like receptors (CLRs), cytosolic RIG-I-like receptors (RLRs), NOD-like receptors (NLRs), and cytosolic DNA receptors (CDRs). Upon activation by PAMPs or DAMPs, PRRs trigger transcription dependent or independent cell events, and subsequently drive the expressions of type I interferons (IFNs), pro-inflammatory cytokines or protease activation, orchestrating innate immune responses (2).

Innate immune DNA receptors include TLR9 and cytosolic DNA receptors (CDRs) (3). Among an array of CDRs, the cyclic GMP-AMP synthase (cGAS) has been the most defined and important DNA receptor (4). It belongs to the nucleotidyltransferase (NTase) family, and can catalyze the substrates ATP and GTP into endogenous cyclic guanosine monophosphate-adenosine monophosphate (2’3’-cGAMP), which, as a second messenger, activates the downstream endoplasmic reticulum (ER) associated adaptor STING (5-8). Activated STING traffics to ER–Golgi intermediate compartment (ERGIC) and Golgi where it recruits TBK1 through the [(D/E)xPxPLR(S/T)D] motif in its C-terminal tail (CTT). The recruited TBK1 is self-transactivated and phosphorylates conserved serine (S) located in the IRF3 recruitment motif (pLxIS) in STING CTT (9). Next, the phosphorylated S serves as a docking site to recruit transcription factor IRF3 which is then phosphorylated by nearby TBK1 (10). The phosphorylated IRF3 forms a dimer and translocates into nucleus to drive the expressions of type I IFNs and the numerous IFN stimulated genes (ISGs), inducing a robust antiviral state (11-13). In addition, STING also activates NF-κB to a low extent via the IKKα/β/γ complex, which is able to synergize with IRF3 in IFN production or drive pro-inflammatory cytokine productions (14).

Beyond the IRF3 and NF-κB mediated signaling, both cGAS and STING are able to activate autophagy including canonical as well as non-canonical autophagy (15). STING triggered autophagy was showed to exert an antiviral role independently of IFNs (16, 17). It was observed that STING is able to mediate ER stress and the unfolded protein response (UPR) through a novel and distinct motif named “the UPR motif” in cyclic dinucleotide binding (CBD) domain, which negatively regulates AKT/TSC/mTOR pathway to enhance canonical autophagy (18, 19). On the other hand, several studies revealed the detailed mechanisms of STING triggered non-canonical autophagy (16, 20-22): Activated STING exits from the ER to the ERGIC and Golgi in a process dependent on the COP-II complex and ARF GTPases, and the ERGIC can serve as membrane sources for LC3 lipidation and autophagosome biogenesis. Different from canonical autophagy, the STING triggered non-canonical autophagy is independent of upstream autophagy regulators such as ULK1, Beclin-1 and Atg9a, but dependent on downstream autophagy regulators such as Atg5 and Atg16L1 (16). The STING induced non-canonical autophagy is mainly initiated by the direct interaction of STING with LC3 through the LC3 interacting regions (LIRs) on STING (21).

Recently, the IFN independent antiviral activity of cGAS-STING pathway has attracted the attentions from researchers (23), and STING induced autophagy may be the contributor to such an antiviral activity (16, 24-26). However, the antiviral activity of STING induced autophagy has not been fully elucidated. In this study, we investigated porcine STING (pSTING) signaling in anti-HSV1 and anti-VSV infections and found that pSTING exerts both IFN and autophagy independent antiviral activity. Further, we also found that STING triggered cell apoptosis may be involved in the resistance from virus infections.

## Materials and methods

### Reagents and antibodies

TRIpure reagent was purchased from Aidlab (Beijing, China). Annexin V-FITC/PI Apoptosis Detection Kit, HiScript® 1st Strand cDNA Synthesis Kit, ChamQ Universal SYBR qPCR Master Mix, 2×Taq Master Mix (Dye plus) and 180 kDa prestained protein marker were all from Vazyme Biotech Co., Ltd (Nanjing, China). The Golden Star T6 Super PCR mix polymerase was from by Tsingke (Nanjing, China). Double-luciferase reporter assay kit was bought from TransGen Biotech (Beijing, China). 2’3’-cGAMP was bought from InvivoGen (Hong Kong, China). Low melting agarose was purchased from Sigma-Aldrich (St. Louis, USA). HSV1-GFP, VSV-GFP and SeV-GFP were preserved in our laboratory. All other chemicals and reagents were analytical grade and obtained commercially.

The mouse anti-Actin mAb and mouse anti-GFP mAb were acquired from Transgen Biotech (Beijing, China). The rabbit STING pAb (19851-1-AP) and rabbit phosphorylated p62 (p-p62) pAb (18420-1-AP) were both purchased from ProteinTech (Wuhan, China). The rabbit TBK1 mAb (3504S), rabbit phosphorylated-TBK1 (p-TBK1, Ser172) mAb (5483S), IRF3 mAb (11904S), HA mAb (3724) and rabbit LC3B (D11) XP^®^ mAb (3868) were all acquired from Cell Signaling Technology (Boston, MA, USA). The rabbit phospho-IRF3 (p-IRF3, Ser385) pAb (MA5-14947) was purchased from Thermo Fisher (Sunnyvale, CA, USA). Rabbit ISG56 pAb was homemade and stored in our lab. HRP goat anti-rabbit IgG (H+L) highly cross-adsorbed secondary antibody and goat anti-mouse IgG (H+L) highly cross-adsorbed secondary antibody were obtained from Sangon Biotech (Shanghai, China).

### Cells and cell transfection

HEK-293T cells (ATCC Cat# CRL-3216) were maintained in Dulbecco modified Eagle medium (DMEM, Hyclone Laboratories, USA) supplemented with 10% fetal bovine serum (FBS, Gibco, USA) and 1% penicillin/streptomycin, and maintained at 37 °C with 5% CO_2_. Porcine macrophages 3D4/21 cells (ATCC Cat# CRL-2843) were cultured in RPMI 1640 medium (Hyclone Laboratories, USA) supplemented with 10% FBS and 1% penicillin/streptomycin, and maintained at 37 °C with 5% CO_2_. Transfection was performed by using the Lipofectamine 2000 (Thermo Fisher, USA) following the manufacturer’s instructions.

### Molecular cloning and gene mutations

The HA tagged pcDNA DEST plasmids of porcine cGAS (pcGAS) and porcine STING (pSTING) were previously constructed, characterized (27, 28) and have been used in our laboratory. The mutation PCR primers of pSTING were designed using QuickChange Primer Design method (https://www.agilent.com) and shown in Supplementary Table 1. The mutation PCR was performed with KOD plus neo polymerase (Toyobo, Shanghai, China) and pcDNA-pSTING-2HA as the template. The PCR products were transformed into competent DMT *E*.*coli* after *Dpn* I digestion, and the resultant mutants were sequence confirmed by DNA sequencing.

### CRISPR gRNA mediated gene knockout (KO) macrophage cell clones

STING^-/-^ and IRF3^-/-^ 3D4/21 cells were previously constructed and used in this study (28). For TBK1 gene knockout, two CRISPR gRNAs were designed using the web tool from Benchling (www.benchling.com), as shown in Supplementary Table 2. The annealed gRNA encoding DNA sequences were cloned into *Bbs*I site of pX458-EGFP and the recombinant pX458-gRNA plasmids were sequence confirmed. The 3D4/21 cells grown in 6-well plates (6-8×10^5^ cells/well) were co-transfected with two pX458-gRNA plasmids using Lipofectamine 2000. Twenty-four hours later, the GFP positive cells were sorted by flow cytometry into 96-well plates for monoclonal cell growth. The individual clones were examined by Western blotting for TBK1 expression and also detected for genomic DNA editing by PCR with primers shown in Supplementary Table 2. The genomic PCR products were cloned into T vector using pClone007 versatile simple vector kit (TsingKe Biological Technology, Beijing, China) and inserted fragments were multiply sequenced. Eight sequences were aligned and analyzed for base insertion and deletion (ins/del) mutations to ensure the availability of TBK1^-/-^ cell clone.

### Promoter-driven luciferase reporter gene assay

293T cells grown in 96-well plates (3×10^4^ cells/well) were co-transfected using Lipofectamine 2000 with ISRE-luciferase reporter or ELAM (NF-κB)-firefly luciferase (Fluc) reporter (10 ng/well) and β-actin Renilla luciferase (Rluc) reporter (0.2 ng/well), together with the indicated plasmids or vector control (5–40 ng/well). The total DNA per well was normalized to 50 ng by adding corresponding empty vectors. Twenty-four hours after transfection, the cells were lysed, and the Fluc and Rluc activities were measured sequentially by double-luciferase reporter assay kit. The results were expressed as fold induction of ISRE or ELAM (NF-κB)-Fluc compared with that of vector control after Fluc normalization by corresponding Rluc.

### RNA extraction, reverse transcription and quantitative PCR (RT-PCR)

Total RNA was extracted using TRIpure reagent following the manufacturer’s suggestions. The extracted RNA was reverse transcribed into cDNA using HiScript 1^st^ strand cDNA synthesis kit, and then the target gene expressions were measured using quantitative PCR with SYBR qPCR master Mix (Vazyme, Nanjing, China) using StepOne Plus equipment (Applied Biosystems). The qPCR program included a denaturation step at 94°C for 30 s, followed by 40 cycles of 94°C for 5 s and 60 °C for 30 s. The relative mRNA levels were normalized to RPL32 mRNA levels, and calculated using 2^−ΔΔCT^ method. The sequences of qPCR primers used were shown in Supplementary Table 3.

### Western blotting analysis

Whole cell proteins were extracted from cells with an RIPA lysis buffer (50 mM Tris pH 7.2, 150 mM NaCl, 1% sodium deoxycholate, 1% Triton X-100). The protein lysate samples were 3:1 mixed with 4×loading buffer and boiled for 5-10 min. The lysate supernatants after centrifugation were run by SDS-PAGE, and then the proteins in gel were transferred to PVDF membrane. The membrane was incubated with 5% skim milk TBST at room temperature (RT) for 1 h, probed with the indicated primary antibodies at 4 °C overnight, washed, and then incubated with secondary antibodies for 1 h at RT. The protein signals were detected by ECL detection substrate and imaging system (Tanon, Shanghai, China).

### Plaque assay

Vero cells were seeded into 24-well plates (3×10^5^ cells/well). Once the cells were grown into monolayer, cells were infected with the tenfold serially diluted cell supernatants from HSV1, VSV or SeV infected cells for 2 h. Then the infected cells were washed with PBS and overlaid by immobilizing medium of 1:1 mixture of warmed 2×DMEM with 4% FBS and a stock solution of heated 1.6% low melting agarose. Plague formations took 4 days for HSV1, and 2 days for VSV and SeV, respectively. Upon completion, the immobilizing medium were discarded by tipping and cells were fixed and stained with crystal violet cell colony staining solution (0.05% w/v crystal violet, 1% formaldehyde, 1×PBS and 1% methanol) for 1 h at RT. Finally, cells were washed with tap water until the clear plaques appeared. The plaques were counted and photos were taken.

### Flow Cytometry for apoptosis detection

The level of cell apoptosis was examined using the Annexin V-FITC/PI Apoptosis Detection Kit. Briefly, the treated cells were harvested by trypsin digestion and washed with the binding buffer, and then re-suspended in the binding buffer. The staining solution of Annexin V-FITC (Fluorescein isothiocyanate isomer I) and PI (propidium iodide) were added successively. The cells were incubated with fluorescein dyes at RT for 15 min in the dark, and the stained cells were immediately detected using flow cytometry. About 10, 000 cell events were collected for analysis of FITC and PI signals detected by channels FL1 and FL3, respectively. The FITC and PI signals were processed with software FlowJo and presented as dot plots.

### Statistical analysis

All the results were the representatives of two or three similar experiments. The data in bar graphs were presented as the mean ± standard deviation (SD) and analyzed using GraphPad Prism 8.0. Student’s *t*-test and *ANOVA* were used for statistical analysis and *P* < 0.05 was considered statistically significant. Statistical signs are indicated as follows: * or ^#^, *P* < 0.05; **, *P* < 0.01; ***, *P* < 0.001 and ns, not significant.

## Results

### pSTING plays a critical role in resisting viral infections

To determine the antiviral effect of porcine cGAS-STING pathway, we transfected porcine (p) cGAS and/or porcine (p) STING into 293T cells followed by infections with HSV1 and VSV, respectively. In HSV1 infected cells, the results showed that pcGAS transfection alone could not induce significant anti-HSV1 effect, while pSTING transfection alone had weak but significant anti-HSV1 effect. The pSTING exhibited more pronounced anti-HSV1 effect upon co-transfection and activation of pcGAS, which were evidenced by GFP fluorescence microscopy, viral GFP Western blotting and viral plaque assay (Fig 1A-D). In the VSV infection, it could be observed that pSTING activated by pcGAS could induce significant anti-VSV effect (Fig 1E-H).

**Figure 1.**
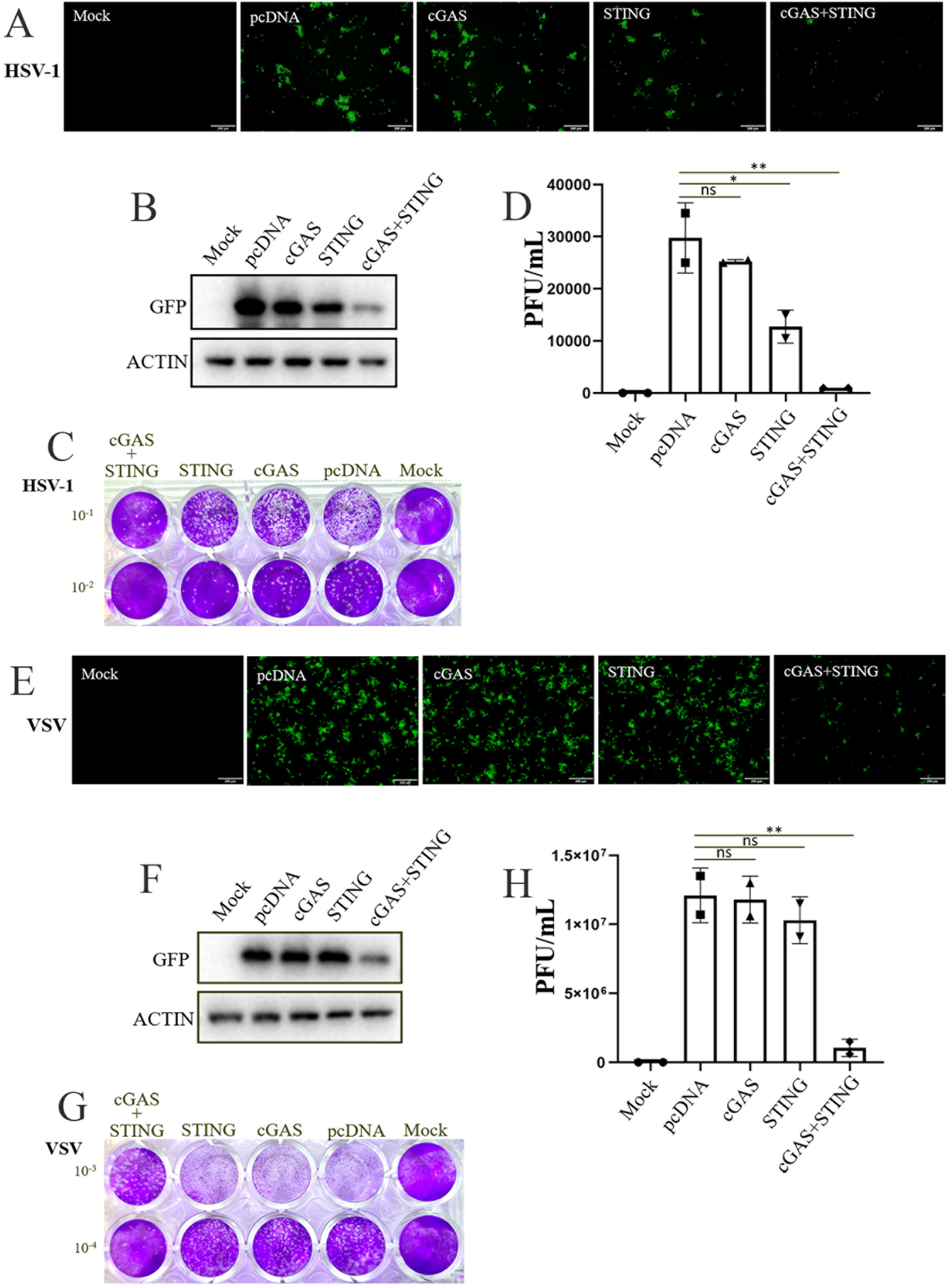
Porcine cGAS-STING signaling restricts HSV1 and VSV infections. (**A**-**D**) 293T cells in 24-well plate (1.5×10^5^ cells/well) were transfected with pcGAS (0.25 μg), pSTING (0.25 μg), pcGAS plus pSTING (0.25 μg + 0.25 μg) or pcDNA control for 24 h. The transfected cells were infected with HSV1-GFP (0.1 MOI) for another 24 h. The viral GFP fluorescence was observed by microscopy (A). The expressions of viral GFP protein were detected by Western blotting (B). HSV1 plaques were showed in (C) and the numbers of plaques were counted and presented in (D). (**E**-**H**) 293T cells in 24-well plate (1.5×10^5^ cells/well) were transfected as in HSV1 infection. The transfected cells were infected with VSV-GFP (0.01 MOI) for another 12 h. The viral GFP fluorescence was observed by microscopy (E). The expressions of viral GFP protein were detected by Western blotting (F). VSV plaques were showed in (G) and the numbers of plaques were counted and presented in (H).

To further verify whether pSTING plays an essential role in antiviral infections, we utilized STING^-/-^ 3D4/21 cells together with normal control 3D4/21 cells stimulated with STING specific agonist 2’3’-cGAMP for infections of HSV1 and VSV, respectively. The results showed that in normal 3D4/21 cells, 2’3’-cGAMP stimulated pSTING signaling exhibited significant anti-HSV1 effect relative to transfection control, whereas in STING^-/-^ 3D4/21 cells, HSV1 replication was upregulated and the 2’3’-cGAMP lost its anti-HSV1 effect (Fig 2A-D). The similar results appeared in VSV infection (Fig 2E-H) Taken together, the results suggested that pSTING plays a key role in antiviral responses.

**Figure 2.**
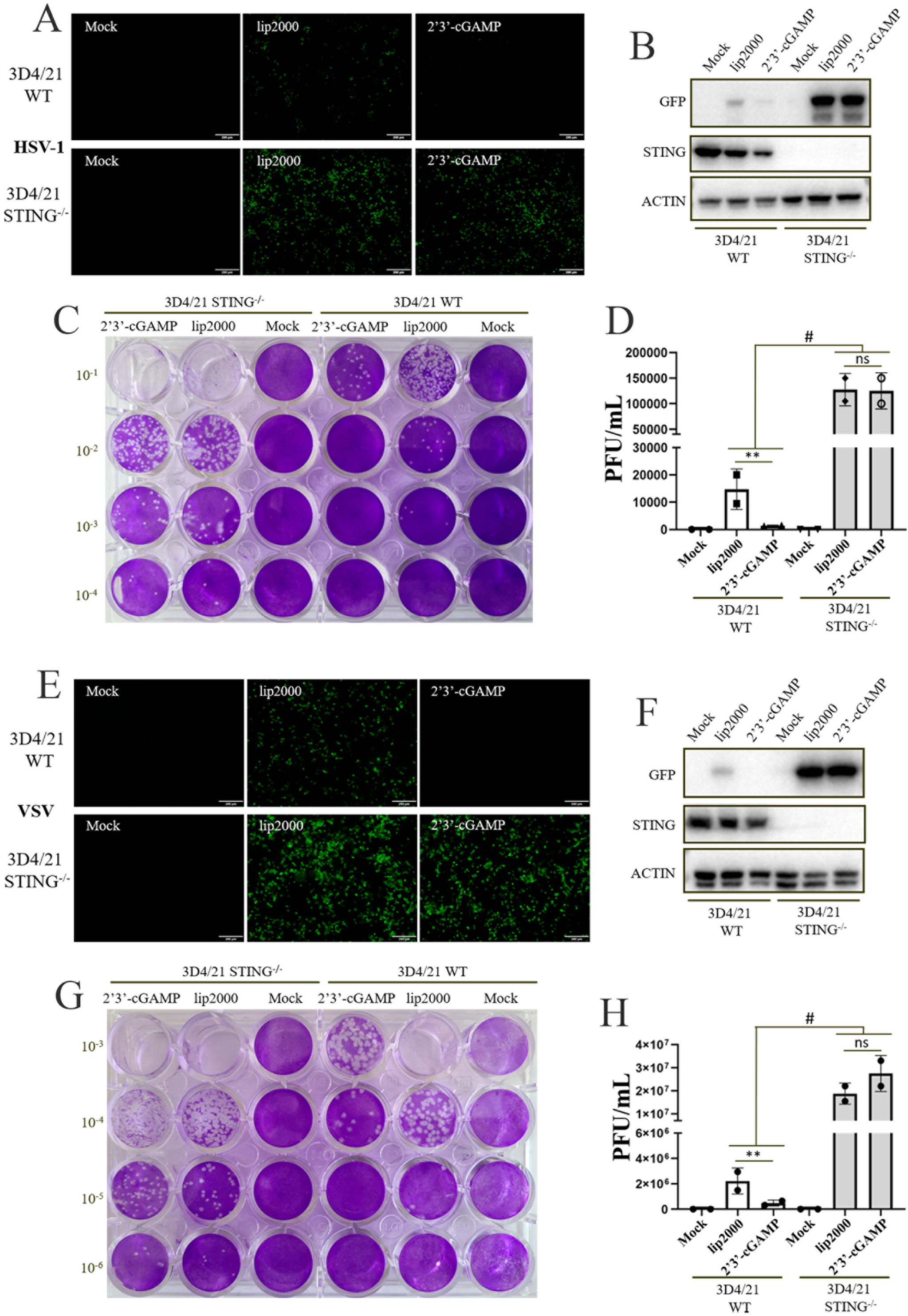
pSTING plays a key role in resisting HSV1 and VSV. (**A**-**D**) STING^-/-^ 3D4/21 and WT 3D4/21 cells grown in 24-well plate (1.5×10^5^ cells/well) were stimulated with transfection of 2’3’-cGAMP (2 μg/mL) by using Lipofectamine 2000. At 12 h post transfection, the stimulated cells were infected with HSV1-GFP (0.1 MOI) for 24 h. The viral GFP fluorescence was observed by microscopy (A) and GFP protein expressions were detected by Western blotting (B). The viral plaques were showed in (C) and the numbers of plaques were plotted in (D). (**E**-**H**) STING^-/-^ 3D4/21 and WT 3D4/21 cells grown in 24-well plate (1.5×10^5^ cells/well) were stimulated as in HSV1 infection. At 12 h post transfection, the stimulated cells were infected with VSV-GFP (0.01 MOI) for 12 h. The viral GFP fluorescence was observed by microscopy (E) and the GFP expressions were detected by Western blotting (F). The viral plaques were showed in (G) and the numbers of plaques were plotted in (H).

### pSTING exerts an antiviral function independently of IFN

The pSTING pLxIS sub is a mutant that has a substituted IRF3 recruitment motif with chicken IRF7 recruitment motif, and no longer induces IFN in mammalian cells including 293T cells (28). We examined the antiviral function of pSTING pLxIS sub in transfected 293T cells. The results showed that the IFN defective pSTING mutant exhibited more pronounced anti-HSV1 and anti-SeV effects compared with WT pSTING (Fig 3A-C, 3G-I and SupFig 1A and C), and comparable anti-VSV effect relative to WT pSTING (Fig 3D-F and SupFig 1B), indicating the IFN independent antiviral function by pSTING.

**Figure 3.**
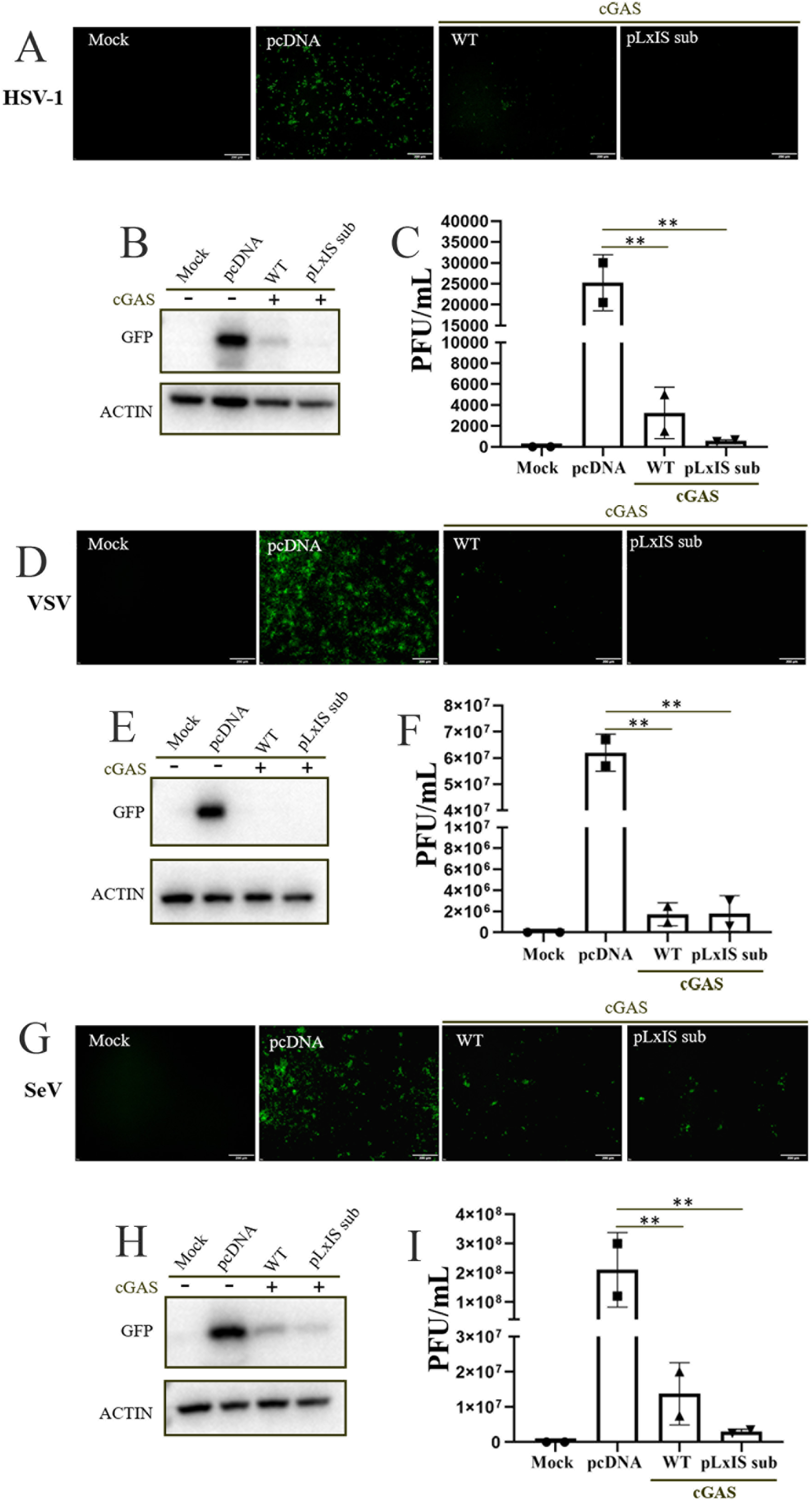
The IFN defective pSTING pLxIS sub exerts antiviral function. (**A**-**C**) 293T cells were transfected with cGAS (0.25 μg) combined with pSTING WT and pLxIS sub (0.25 μg each) for 24 h, with pcDNA transfection and mock as controls. The cells were next infected with HSV1-GFP (0.1 MOI) for another 24 h, followed by detection of viral replications with GFP fluorescence (A), GFP protein expression (B) and viral plaque assay (C). (**D**-**I**) 293T cells were transfected as in HSV1 infection, and infected with VSV-GFP (0.01 MOI, D-F) for 12 h and SeV-GFP (0.01 MOI, G-I) for 12 h. The viral replications were detected by GFP fluorescence transfection (D and G), GFP protein expressions (E and H) and plaque assay (F and I).

In order to further validate the IFN independent antiviral function of pSTING, we constructed two additional pSTING mutants S365A and ΔCTT. The serine 365 of pSTING is the TBK1 phosphorylation site and the phosphorylated serine 365 is for IRF3 recruitment and downstream IFN induction. The pSTING C-terminal tail (CTT) contains both TBK1 recruitment motif and IRF3 recruitment motif, and is thus essential for IFN induction. As expected, both pSTING mutants S365A and ΔCTT lost IFN activity in ISRE promoter assay (Fig 4A). Nevertheless, both mutants kept NF-κB activity with S365A even harboring significantly higher NF-κB activity than WT pSTING indicating the competition between IRF3 and NF-κB signals (Fig 4B). Consistently, RT-qPCR assay showed that pSTING S365A and ΔCTT could not induce IFN-β, ISG56 and ISG60, but could induce IL-8 and TNF-α (Fig 4C). In Western blotting, pSTING S365A could induce the TBK1 phosphorylation, but not the IRF3 phosphorylation and downstream ISG56 production, whereas pSTING ΔCTT could not induce the TBK1 and IRF3 phosphorylations, and ISG56 production (Fig 4D). The antiviral activity of pSTING S365A and ΔCTT were examined in transfected 293T cells and the results showed that both mutants S365A and ΔCTT still possessed the anti-HSV1 activity (Figure 4E-G and SupFig 1D) and anti-VSV activity (Figure 4H-J and SupFig 1E) despite of the defects in IFN activity. Taken together, these results clearly suggested that pSTING plays a strong antiviral role independently of IFN.

**Figure 4.**
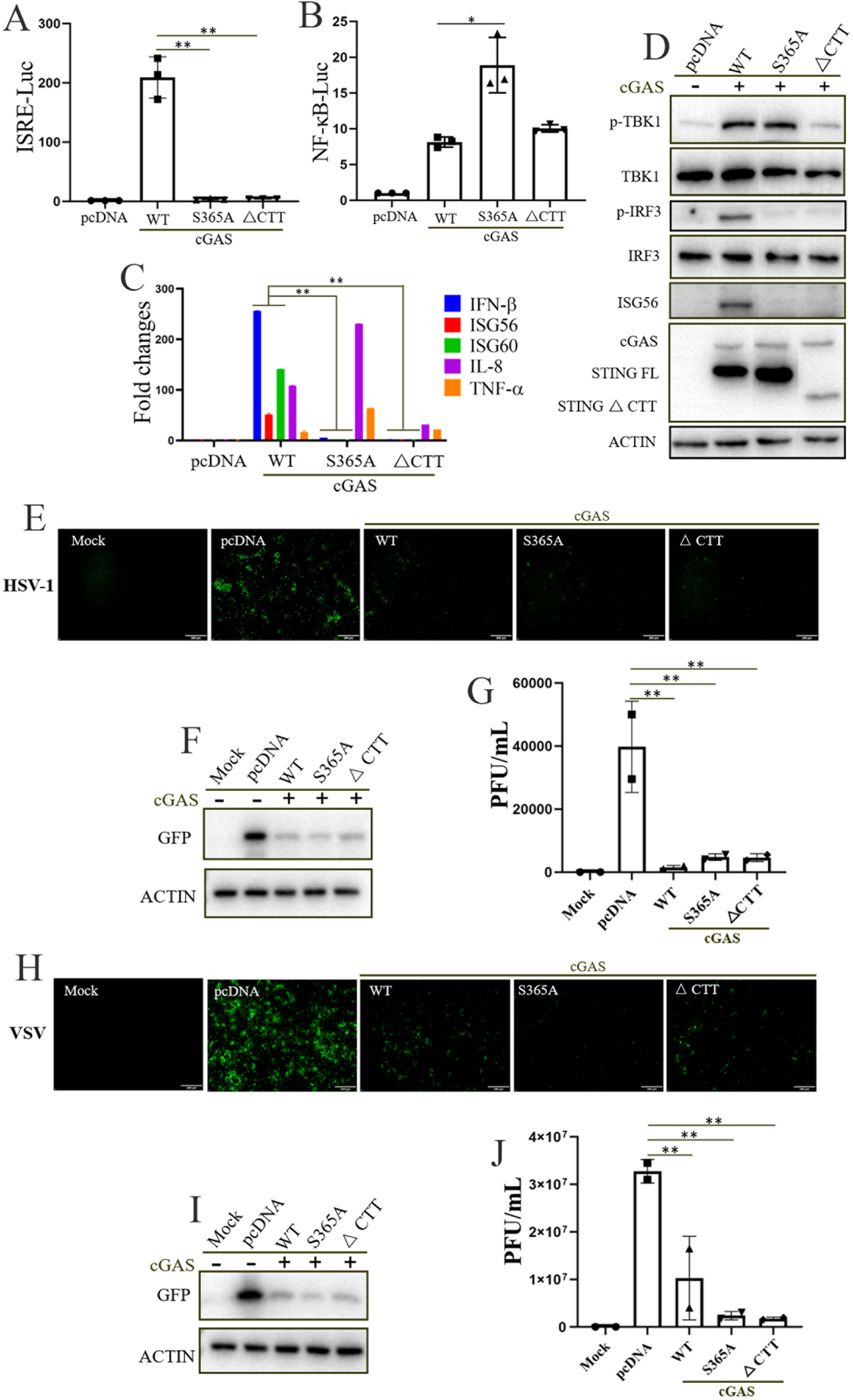
The IFN defective pSTING S365A and ΔCTT also exert antiviral function. (**A-B**) 293T cells in 96-well plate (3×10^4^ cells/well) were transfected with the indicated combinations of pcGAS (20 ng) and pSTING WT/S365A/ΔCTT (20 ng each), or empty vector (40 ng), together with ISRE Fluc (10 ng) (A) or NF-κB Fluc (10 ng) (B) plus Rluc (0.2 ng). Luciferase activities were measured 24 h after transfection. (**C**) 293T cells grown in 12-well plate (3×10^5^ cells/well) were transfected with the indicated combinations of pcGAS (0.5 μg) and pSTING WT/S365A/ΔCTT (0.5 μg each) for 48 h. The RNA expressions of hIFN-β, hISG56, hISG60, hIL-8 and hTNF-α were analyzed by RT-qPCR. (**D**) 293T cells grown in 24-well plate (1.5×10^5^ cells/well) were transfected with the indicated combinations of pcGAS (0.25 μg) and pSTING WT/S365A/ΔCTT (0.25 μg each) or empty pcDNA vector (0.5 μg) for 24 h, and the expressions of p-TBK1, TBK1, p-IRF3, IRF3, ISG56, cGAS-HA and STING-HA were examined by Western blotting. (**E-J**) 293T cells were transfected with pcGAS (0.25 μg) combined with pSTING WT/S365A/ΔCTT (0.25 μg each) for 24 h, with the pcDNA transfection and mock as controls. The treated cells were infected with HSV1-GFP (0.1 MOI, E-G) for 24 h and VSV-GFP (0.01 MOI, H-J) for 12 h, followed by detection of viral replications with GFP fluorescence (E and H), GFP protein expressions (F and I) and plaque assay (G and J).

### pSTING induced autophagy does not depend on IFN but affects IFN

STING can not only induce IFN and inflammatory cytokines, but also induce autophagy (16, 20, 21). The autophagy inductions by pSTING and three IFN defective pSTING were examined in transfected 293T cells, and the results showed that three IFN defective pSTING mutants pLxIS sub, S365A and ΔCTT all induced comparable LC3 lipidation (LC3-II) and p62 phosporylation (p-p62) relative to WT pSTING, suggesting that pSTING induced autophagy was independent of IFN (Fig 5A). Previously, a small region of residues 330–334 of human STING was identified as important for autophagy induction and the point mutations of L333 and R334 within this region of STING abrogated LC3 lipidation (11). We made the equivalent point mutations in pSTING WT, S365A and ΔCTT, respectively, and tested the autophagy induction by these L333A/R334A mutants in transfected 293T cells. Compared with WT pSTING, the pSTING L333A/R334A had decreased levels of LC3-II (lane 12 vs lane 9, Fig 5E). Similarly, for both pSTING S365A and ΔCTT, the L333A/R334A mutation also decreased the levels of LC3-II and p-p62 (Fig 5B). Because mutation L333A/R334A also decreased the expressions of various pSTING proteins (Fig 5B and E), the pSTING WT, S365A and ΔCTT protein expressions were titrated and adjusted for equal expressions as the corresponding L333A/R334A mutants (Fig 5C, D and F). After avoiding the disturbance of pSTING expressions, the L333A/R334A mutation was validated to dampen the autophagy by pSTING WT, S365A and ΔCTT, respectively (Fig 5C, D and F).

**Figure 5.**
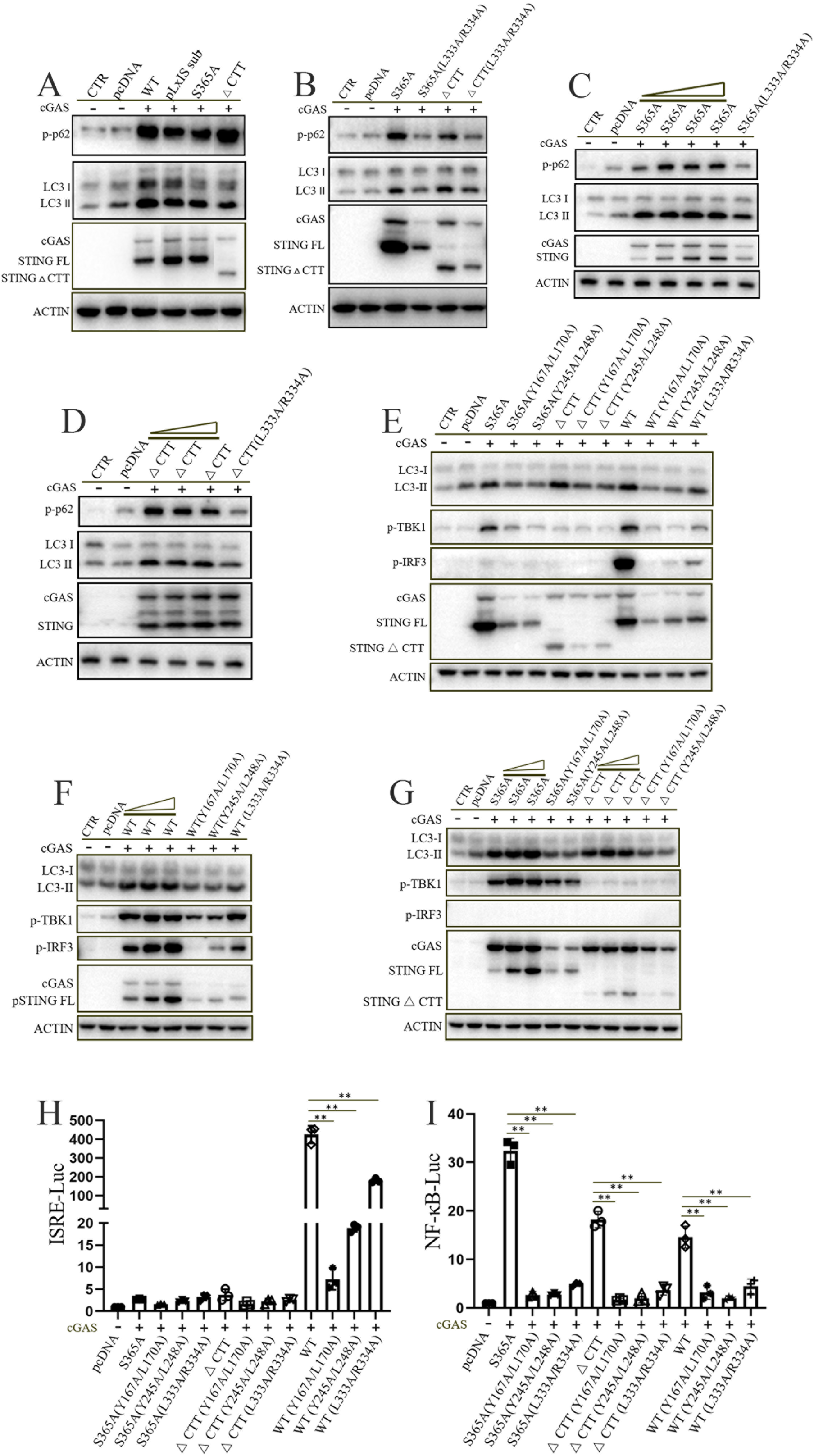
The analysis of autophagy, IFN and NF-κB activities of various pSTING mutants. (**A**) 293T cells grown in 24-well plate (1.5×10^5^ cells/well) were transfected with the indicated combinations of pcGAS (0.25 μg) and pSTING WT/pLxIS sub/S365A/ΔCTT (0.25 μg each) for 24 h, and the indicated protein expressions were analyzed by Western blotting. (**B**) 293T cells were transfected with the indicated combinations of pcGAS (0.25 μg) plus pSTING S365A, ΔCTT and the corresponding L333A/R334A mutants (0.25 μg each) for 24 h, followed by Western blotting analysis. (**C** and **D**) 293T cells were transfected with the indicated combinations of pcGAS (0.25 μg) plus S365A gradient (0.1, 0.15, 0.2, 0.25 μg) and S365A L333A/R334A (0.25 μg) (C), ΔCTT gradient (0.05, 0.15, 0.25 μg) and ΔCTT L333A/R334A (0.25 μg) (D) for 24 h. (**E**) 293T cells were transfected with the pcGAS (0.25 μg) combined with pSTING WT, S365A, ΔCTT, various Y167A/L170A, Y245A/L248A and L333A/R334A mutants (0.25 μg each), as indicated, for 24 h. (**F**) 293T cells were transfected with pcGAS (0.25 μg) combined with pSTING WT gradient (0.05, 0.15, 0.25 μg) and the mutants Y167A/L170A, Y245A/L248A and L333A/R334A (0.25 μg each), for 24 h. (**G**) 293T cells were transfected with pcGAS (0.25 μg) combined with pSTING S365 gradient (0.05, 0.15, 0.25 μg), ΔCTT gradient (0.05, 0.15, 0.25 μg) and the corresponding mutants Y167A/L170A, Y245A/L248A (0.25 μg each), for 24 h. (**H and I**) 293T cells in 96-well plate were transfected with pcGAS (20 ng) combined with pSTING WT, S365A, ΔCTT and various mutants as indicated (20 ng each), plus ISRE FLuc and Rluc (H), or plus ELAM (NF-κB) Fluc and Rluc (I). Luciferase activities were measured 24 h post transfection.

Human STING was showed to directly activate autophagy by interacting with LC3 through the LC3 interacting regions (LIRs) involving in LIR4, LIR6 and LIR7 of STING CBD domain (21). Based on the alignment of human and porcine STING (Sup Fig 2), only LIR4 and LIR7 are present in pSTING and point mutations of LIR4 (Y167A/L170A) and LIR7 (Y245A/L248A) were made on pSTING WT, S365A and ΔCTT, respectively and separately (Fig 5E). The pSTING mutants Y167A/L170A and Y245A/L248A were tested for autophagy induction in transfected 293T cells and the results showed that either Y167A/L170A or Y245A/L248A mutation in various pSTINGs decreased the LC3-II levels to the basal level as in pcDNA transfected control (Fig 5E). Again, after adjusting the protein expressions of pSTING WT, S365A and ΔCTT as low as the corresponding Y167A/L170A or Y245A/L248A mutants (Fig 5F and G), the mutation Y167A/L170A or Y245A/L248A was validated to abrogate the autophagy by pSTING WT, S365A and ΔCTT, respectively (Fig 5E-G).

The above pSTING autophagy mutants were further examined for IFN and NF-κB activity. L333A/R334A, Y167A/L170A and Y245A/L248A mutations all decreased pSTING induced levels of p-TBK1 and p-IRF3 (lanes 9-12, Fig 5E) as well as ISRE promoter activity (Fig 5H). On the other hand, all these autophagy mutations also decreased the NF-κB promoter activity triggered by pSTING WT, S365A and ΔCTT, respectively (Fig 5I). Taken together, the above results suggest that pSTING triggers IFN independent autophagy; however, pSTING autophagy influences its IFN as well as NF-κB activity.

### pSTING enables resistance to viral infections independently of both IFN and autophagy

Since it seems that the antiviral function of pSTING is associated with its autophagy activity, the various pSTING autophagy mutants in the context of pSTING WT, S365A and ΔCTT were tested for the anti-HSV1 and anti-VSV effects in transfected 293T cells. The antiviral effect was indicated and reflected by GFP microscopy (Fig 6A and F), GFP Western blotting (Fig 6B and G) and plaque assay (Fig 6C-E and H-J). Surprisingly, the various Y167A/L170A and Y245A/L248A mutants of pSTING WT, S365A and ΔCTT, in which either the mutation abrogated the autophagy activity, exhibited comparable anti-HSV1 effect relative to corresponding pSTING WT, S365A and ΔCTT (Fig 6A-E and SupFig 3A-C). On the contrary, the various L333A/R334A mutants of pSTING WT, S365A and ΔCTT, in which the mutation dampened the autophagy activity, exhibited strong and significant anti-HSV1 effect despite of the significant decreased antiviral activity relative to the corresponding pSTING WT, S365A and ΔCTT (Fig 6A-E and SupFig 3A-C). Similar results were obtained in anti-VSV infection setting (Fig 6F-J and SupFig 3D-F).

**Figure 6.**
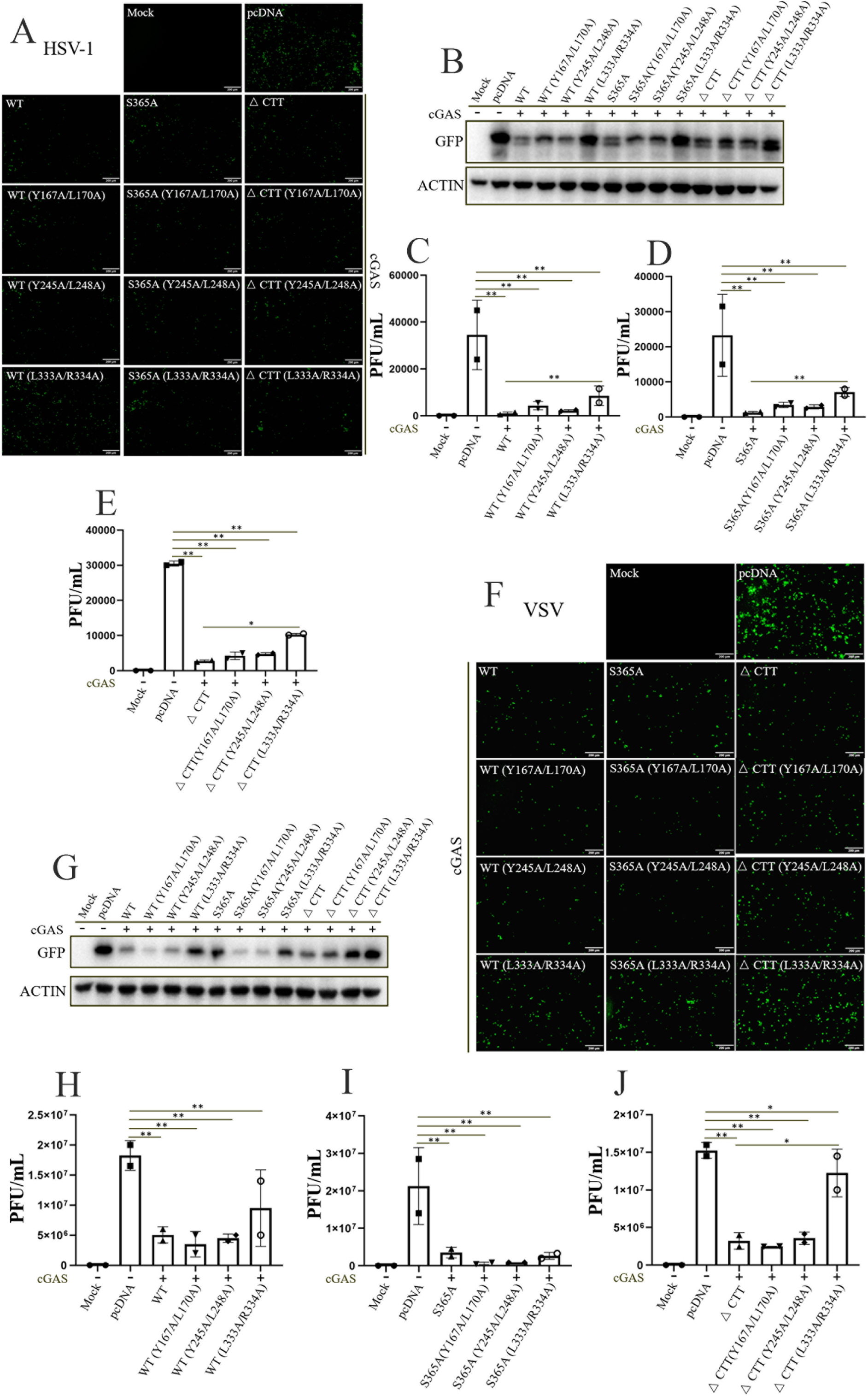
pSTING can resist viruses independently of both IFN and autophagy activity. (**A-E**) 293T cells were transfected with pcGAS (0.25 μg) combined with pSTING WT, S365A, ΔCTT, various mutants and pcDNA control as indicated (0.25 μg each) for 24 h. The transfected cells were infected with HSV1-GFP (0.1 MOI) for another 24 h, followed by detection of viral replications with GFP fluorescence (A), GFP protein expressions (B) and plaque assay (C-E). (**F-J**) 293T cells were transfected as in HSV1 infection, and the transfected cells were infected with VSV-GFP (0.01 MOI) for 12 h, followed by detection of viral replications with GFP fluorescence (F), GFP protein expressions (G) and plaque assay (H-J).

These results clearly suggest that pSTING exerts an antiviral function in a way not only independent of its IFN activity, but also independent of its autophagy activity. It also further suggests that pSTING exerts an antiviral function in a way independent of both IFN and autophagy.

### pSTING induces apoptosis independently of both IFN and autophagy

STING has been implicated to induce cell apoptosis (29). Thus, we continued to explore whether pSTING can induce apoptosis after losing interferon and autophagy activities. To this end, pSTING WT, pLxIS sub, S365A, ΔCTT and all autophagy mutants in the context of pSTING WT, S365A, ΔCTT were transfected into 293T cells to detect their apoptotic activity (Fig 7). Flow cytometry showed that all pSTING including those defective of both IFN and autophagy were able to induce significant apoptosis (Fig 7A-C). Among various pSTING mutants, the L333A/R334A mutants of pSTING WT, S365A and ΔCTT exhibited significant decreased apoptosis relative to the corresponding pSTING WT, S365A and ΔCTT, but still kept the ability to induce both early and late cell apoptosis (Fig 7B and 7C). Since pSTING induces apoptosis as well as antiviral effects independently of both IFN and autophagy, it indicates that pSTING induced apoptosis and antiviral activity are associated, and the induction of apoptosis may be responsible for the pSTING antiviral function.

**Figure 7.**
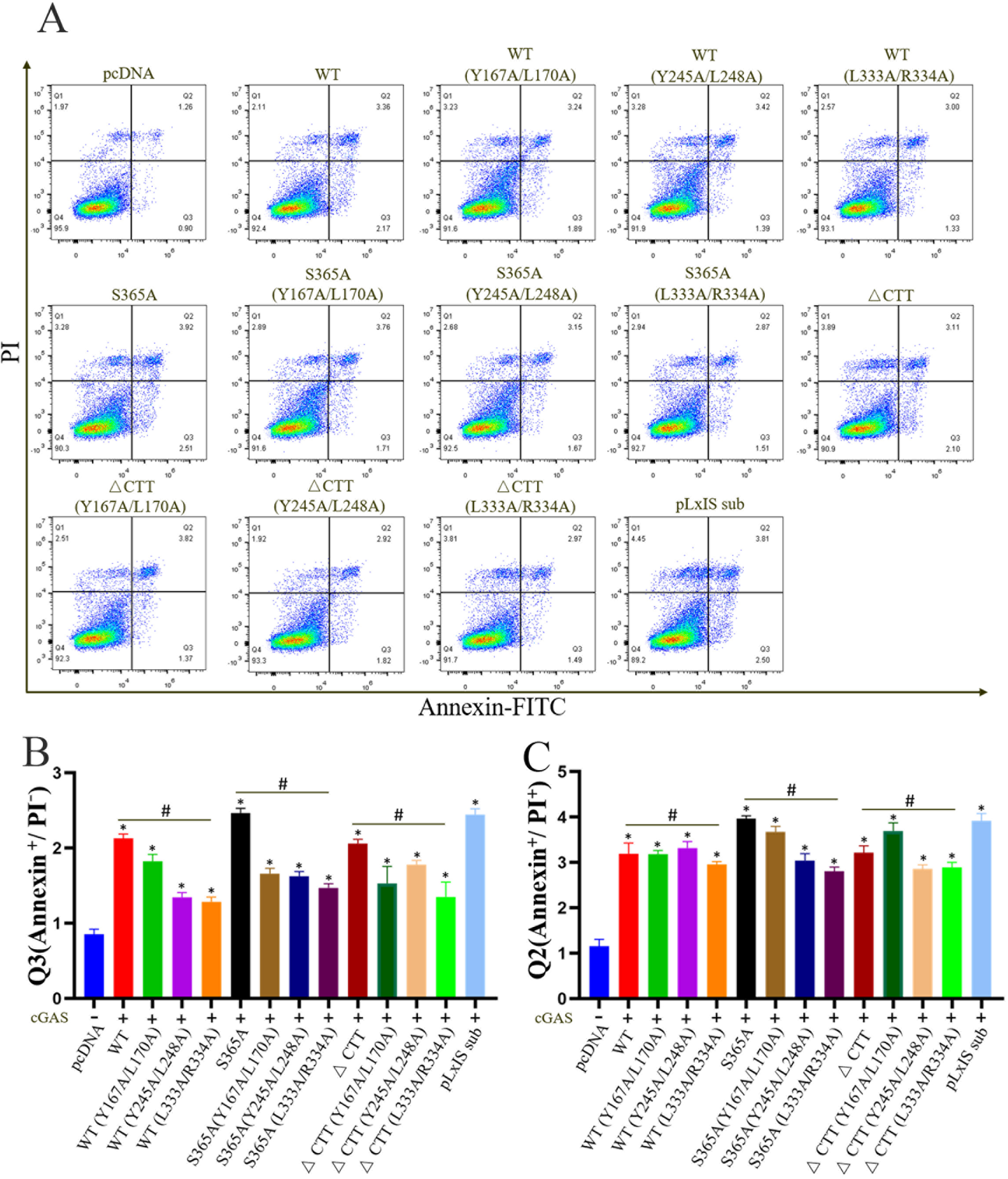
pSTING can induce apoptosis independently of both IFN and autophagy activity. (**A-C**) 293T cells were transfected with pcGAS (0.25 μg) combined with pSTING WT, S365A, ΔCTT, various mutants and pLxIS sub, as indicated (0.25 μg each) for 48 h. The apoptosis of transfected cells was analyzed using Annexin V-FITC/PI staining followed by flow cytometry. The dot plots of cell apoptosis in various samples were presented in A. The early apoptosis as indicated by Q3 (Annexin^+^/PI^+^) and late apoptosis as indicated by Q2 (Annexin^+^/PI^-^) were plotted as bar graphs in B and C, respectively.

### Both TBK1 and IRF3 are necessary for pSTING to exert antiviral function

Next, we explored the roles of downstream TBK1 and IRF3 in the antiviral functions of pSTING. To this end, TBK1 and IRF3 were knocked out from porcine macrophages 3D4/21, respectively, through CRISPR-Cas9 approach (Fig 8). In TBK1^-/-^ 3D4/21 cells, both HSV1 and VSV replications were significantly upregulated and 2’3’-cGAMP activation of pSTING lost the anti-HSV1 (Fig 8A-C) and anti-VSV (Fig 8D-F) effects relative to normal 3D4/21 cells, as evidenced by GFP microscopy, GFP Western blotting and plaque assay (Fig 8A-F and SupFig 4A-B). In IRF3^-/-^ 3D4/21 cells, the same viral infection results were obtained, i.e. both HSV1 and VSV replications were significantly upregulated and 2’3’-cGAMP lost the antiviral function (Fig 8G-L and SupFig 4C-D). These results suggest that although the antiviral function of pSTING is independent of the activation of signal axis TBK1-IRF3 and subsequent production of downstream IFN, the presence of both TBK1 and IRF3 are required.

**Figure 8.**
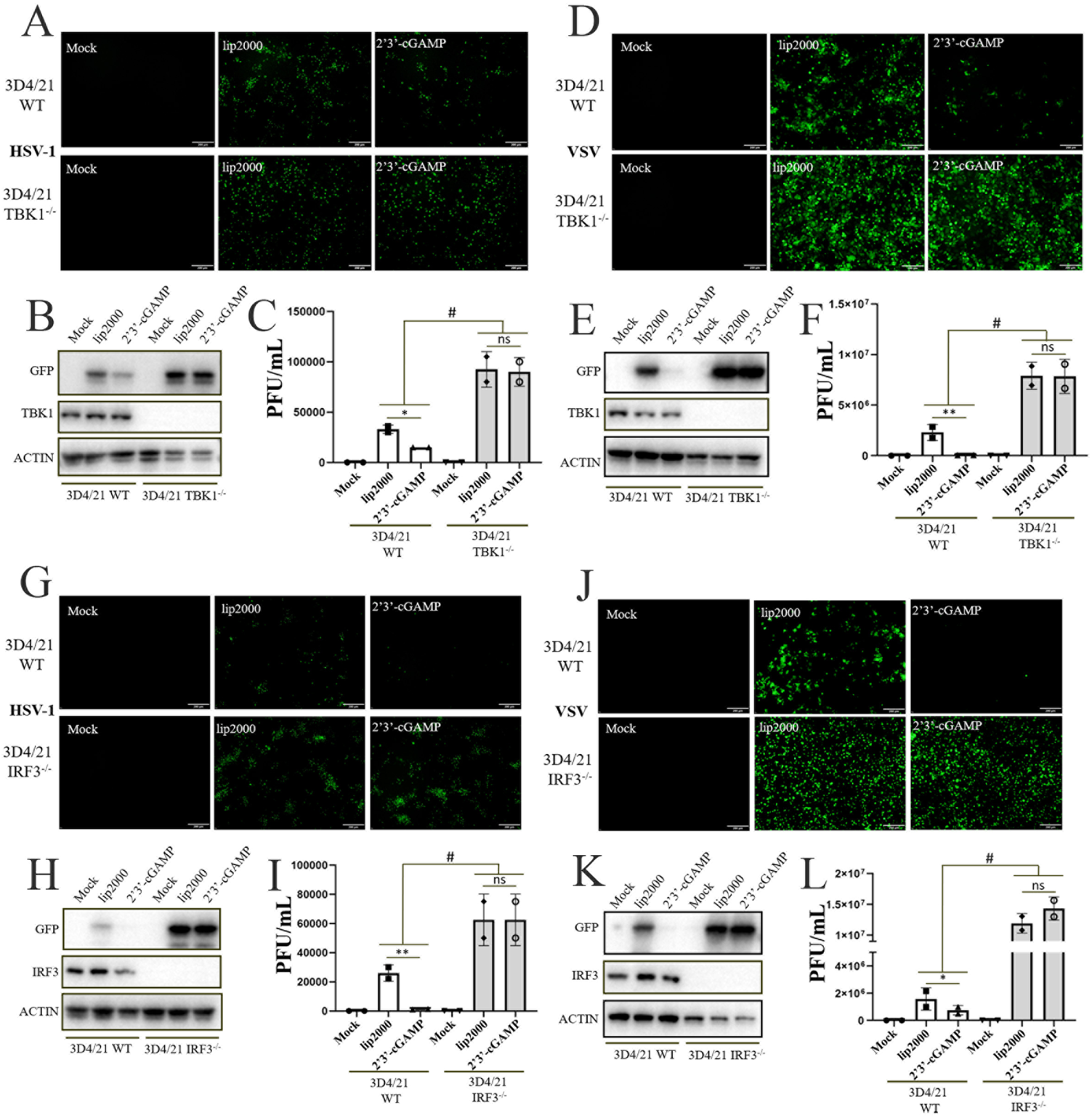
The presences of TBK1 and IRF3 are necessary for the antiviral function of pSTING. (**A**-**F**) TBK1^-/-^ 3D4/21 and WT 3D4/21 cells in 24-well plate were stimulated with transfection of 2’3’-cGAMP (2 μg/mL) using Lipofectamine 2000 for 12 h. The stimulated cells were infected with HSV1-GFP (0.1 MOI, A-C) for 24 h and VSV-GFP (0.01 MOI, D-F) for 12 h, respectively, followed by detection of viral replications with GFP fluorescence (A and D), GFP protein expressions (B and E), and plaque assay (C and F). (**G**-**K**) IRF3^-/-^ 3D4/21 and WT 3D4/21 cells in 24-well plate were stimulated with transfection of 2’3’-cGAMP (2 μg/mL) for 12 h. The stimulated cells were infected with HSV1-GFP (0.1 MOI, G-I) for 24 h and VSV-GFP (0.01 MOI, J-L) for 12 h, respectively, followed by detection of viral replications with GFP fluorescence (G and J), GFP protein expressions (H and K), and plaque assay (I and L).

## Discussion

The cGAS-STING pathway mediated IFN signaling plays a key role in antiviral infections (13). However, it has been appreciated that the cGAS-STING mediated IFN independent signal events also play an important role in restricting HSV1 infection (17, 30, 31); nevertheless, the nature of IFN-independent antiviral activity of STING remains unclear (23). Our study first confirmed the IFN independent antiviral activity of pSTING (Fig 3 and Fig 4) and next explored the nature of such an IFN independent antiviral activity. In addition to IFN production, STING signaling at least activates autophagy, NF-κB and cell death (23, 32). Firstly, despite several studies suggested that STING induced autophagy mediates antiviral function (16, 24-26), our study clearly demonstrated that STING induced autophagy is dispensable for either anti-HSV1 or anti-VSV activity. Secondly, two other studies suggested that NF-κB is implicated in STING antiviral activity (31, 33). In our study, the autophagy defective pSTING mutants also showed a substantial reduction of NF-κB activity (Fig 5I), but the antiviral activity of these pSTING mutants was not altered (Fig 6), indicating NF-κB is also dispensable for STING antiviral activity. Finally, we found that pSTING can induce cell apoptosis independently of IFN and autophagy and the induced apoptosis is clearly associated with the antiviral activity (Fig 7). Therefore, our study revealed the fancy machinery utilized by STING to restrict and resist viral infections.

In this study, we confirmed that pSTING induces IFN independent autophagy, and both LIR4/7 and L333/R334 located in CBD of pSTING are essential for autophagy induction. Among these sites, LIR4/7 are no question involved in STING induced non-canonical autophagy by mediating the interaction of STING and LC3 (21). Although the residues L333/R334 and the corresponding small region (residues 330–334) of human STING were identified as important for autophagy, the machinery of these STING sites inducing non-canonical autophagy is unknown (16). We noticed that this small region is located within the STING UPR motif (residues 322–343), which was initially identified as responsible for STING activated ER stress and subsequent T cell death (19). The STING induced ER stress likely leads to canonical autophagy with mTOR inactivation involved (18). Therefore, LIR4/7 and L333/R334 seem participate in the induction of autophagy with distinct machinery; nevertheless, both sites are implicated in the IFN and NF-κB signals of pSTING (Fig 5H and 5I) and neither is required for antiviral function of pSTING (Fig 6).

STING weakly induces NF-κB signaling relative to IRF3 signaling, but the exact mechanism remains unresolved (32). On the one hand, TBK1 was showed to be required for or participate redundantly with IKK? in the STING induced NF-κB activation (14, 34, 35). On the other hand, STING kept the ability to activate NF-κB without the CTT domain, which excluded the participation and requirement of TBK1 (36, 37). Our results showed that pSTING ΔCTT is able to induce comparable NF-κB (Fig 4B and Fig 5I), suggesting CTT and TBK1 are not necessary for STING activated NF-κB signaling that is similar case for autophagy. Additionally, pSTING induced NF-κB activity was positively regulated by autophagy (Fig 5I), indicating both may be intrinsically linked. The results are favored by the invertebrate STINGs which lack the CTT, but enable activation of NF-κB and autophagy and antiviral functions (16, 38). It is possible that STING may activate NF-κB, alternatively, depending on the signal in CBD of oligomerized STING (32). In terms of the antiviral function, our results indicate that NF-κB is also not necessary for STING antiviral activity.

STING contributes to different types of cell death, such as apoptosis, lysosomal cell death (LCD), pyroptosis, necroptosis and so on (32, 39). Our study found that pSTING can induce cell apoptosis independently of its IFN and autophagy activities, and further that the induced apoptosis is well associated with the antiviral activity (Fig 6 and 7). These results suggest that STING activated apoptosis might exert antiviral function even though it has been shown to induce B cell/T cell death and immune evasion (19, 29), but never direct and clear antiviral functions. STING was shown to induce cell apoptosis via its UPR motif activating ER stress or via downstream IRF3 binding with Bax (39). Our study also showed that the presence of both TBK1 and IRF3 are required for STING induced antiviral activity (Fig 8). The results suggest that both TBK1 and IRF3 might participate in STING induced cell apoptosis and antiviral function, but in a way independent of IFN.

In summary, our study reveals that pSTING exerts an intriguing antiviral function independent of both IFN and autophagy (SupFig 5). The STING antiviral function may be mediated by cell apoptosis, or by several STING signaling events including IFN, NF-κB, autophagy and/or apoptosis, likely in a redundant manner. Considering the conservation of cGAS-STING pathway along evolution, our findings thus provide deep insights into the STING immune biology in viral infections.

## Abbreviations used in this article

cGAS,: cyclic GMP-AMP synthase;
2’3’-cGAMP,: cyclic guanosine monophosphate-adenosine monophosphate;
STING,: stimulator of interferon gene;
TBK1,: TANK-binding kinase 1;
IFN,: interferon;
IRF,: IFN regulatory factor;
ISG,: IFN-stimulated gene;
NF-κB,: nuclear factor kappa light chain enhancer of activated B cells;
poly dA:dT,: poly deoxyadenylic-deoxythymidylic acid;
HSV-1,: herpes simplex virus 1;
VSV,: vesicular stomatitis virus;
SeV,: sendai virus.

## Author contribution statement

S.J, N.X, J.L, Y.Z, Q.C, JJ.Z and Y.W performed the experiments; W.Z, N.C, and F.M contributed to data analysis; S.J and JZ.Z conceived the idea and wrote the paper.

## Acknowledgments

The work was partly supported by the National Natural Science Foundation of China (32172867; 31872450), and A Project Funded by the Priority Academic Program Development of Jiangsu Higher Education Institutions (PAPD). S.J was supported by the Postgraduate Research & Practice Innovation Program of Jiangsu Province(Yangzhou University)(KYCX21_3272).

## Data Availability Statement

The raw data presented in this study are all available upon request.

## Figure legends

**Supplemental Figure 1 to Fig 3 and 4**. Antiviral activity of IFN defective pSTING mutants by plaque assay (**A-C**) The plaque assay showing the activity of anti-HSV1 (A), anti-VSV (B) and anti-SeV (C) by pSTING WT and pLxIS sub in Fig 3. (**D** and **E**) The plaque assay showing the activity of anti-HSV1 (D) and anti-VSV (E) by pSTING WT, S365A and ΔCTT in Fig 4.

**Supplemental Figure 2**. The alignment of STING amino acid sequences from chicken, mouse, human and pig. The three regions of STING including N terminal four transmembrane (TM), middle cyclic dinucleotide-binding domain (CBD) and C terminal tail (CTT) are marked with different colors underneath. The IRF3 binding motif and TBK1 binding motif are blue boxed. The UPR motif and LIR4, 6, 7 motifs are red boxed. In UPR motif, the key residues L333/R334 are green boxed, whereas in LIR motifs, the conserved key residues Y and L are marked with underlying stars. Based on the conserved key residues Y and L, porcine STING have LIR4 and 7 motifs but not LIR6 motif.

**Supplemental Figure 3 to Fig 6**. Antiviral activity of various autophagy defective pSTING mutants by plaque assay. (**A**-**C**) The plaque assay showing the anti-HSV1 activity by pSTING WT and the related autophagy mutants (A), S365A and the related autophagy mutants (B), ΔCTT and the related autophagy mutants (C). (**D**-**F**) The plaque assay showing the anti-VSV activity by pSTING WT and the related autophagy mutants (D), S365A and the related autophagy mutants (E), ΔCTT and the related autophagy mutants (F).

**Supplemental Figure 4 to Fig 8**. Viral replication in TBK1^-/-^ and IRF3^-/-^ 3D4/21 reflected by plaque assay. (**A** and **B**) The plaque assay showing the HSV1 replication (A) and VSV replication (B) in TBK1^-/-^ 3D4/21 and WT 3D4/21 cells with or without 2’3’-cGAMP stimulation. (**C** and **D**) The plaque assay showing the HSV1 replication (C) and VSV replication (D) in IRF3^-/-^ 3D4/21 and WT 3D4/21 cells with or without 2’3’-cGAMP stimulation.

**Supplemental Figure 5**. The schematic diagram of pSTING antiviral activity. It is well known that STING exerts antiviral function through the action of downstream type I IFN, and it was also suggested that STING induced NF-κB and autophagy mediate antiviral activity. However, in our study, pSTING exerts antiviral function independent of its IFN, autophagy as well as NF-κB activity. Instead, the antiviral activity of pSTING is well associated with its apoptosis activity, indicating the apoptosis activity of pSTING may mediate its antiviral function. Alternatively, multiple signaling events of pSTING mediate the antiviral function, together and redundantly.

